## Supplement for "Agent-based model predicts that layered structure and 3D movement work synergistically to reduce bacterial load in 3D in vitro models of tuberculosis granuloma"

Supplementary Information

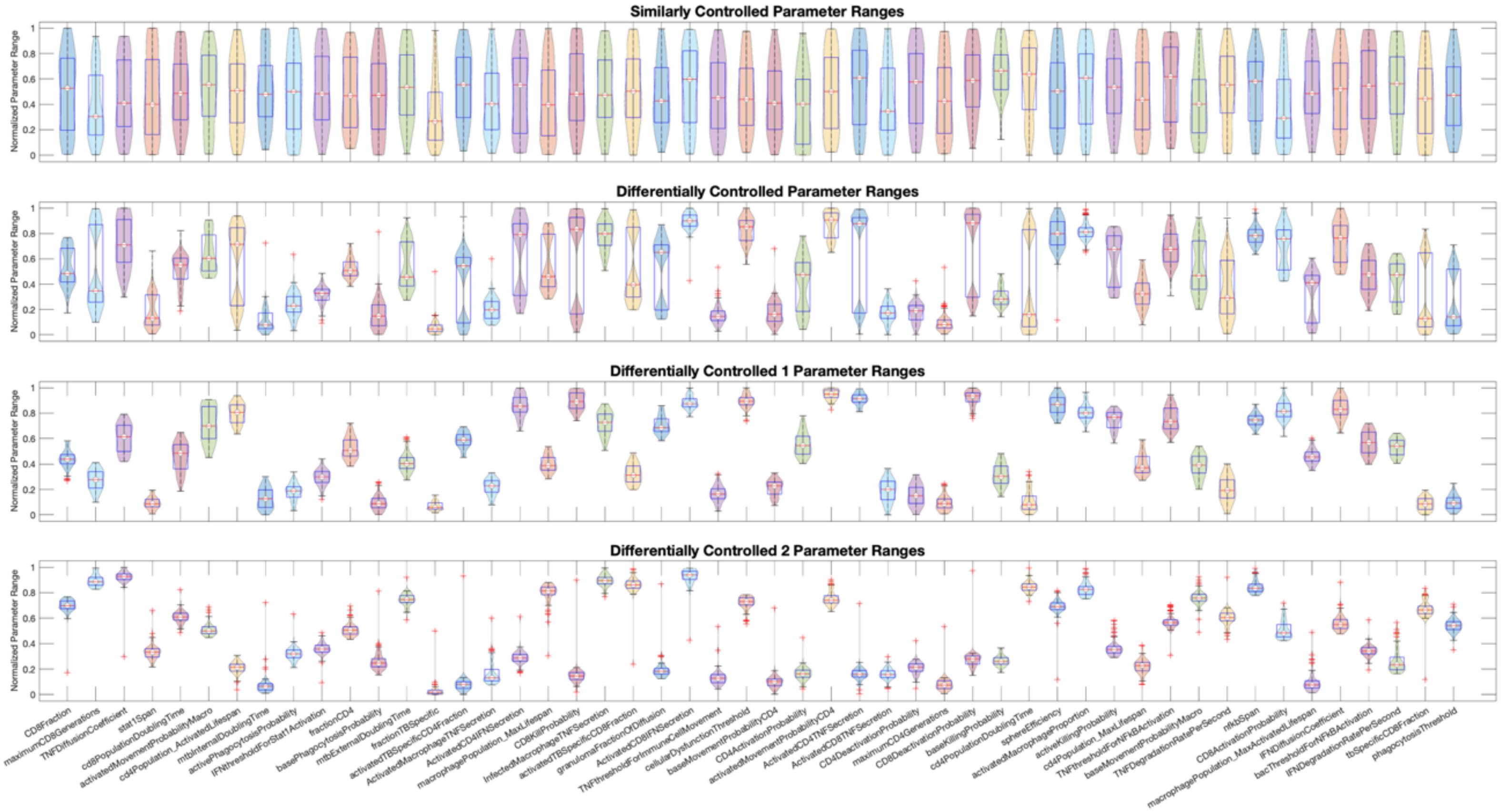


**Figure 1:** The distributions of parameters for similarly controlled simulations, differentially controlled simulations, DC-1 simulations, and DC-2 simulations. Each parameter's range has been normalized from 0 to 1 based on the initial lower and upper bounds of the parameters ranges as given in **Table 1**.

**Table 1:** Parameters that are varied during calibration. Initial ranges are either determined by literature, estimated through preliminary simulations (e), or broadened to the full mathematically possible range (f). The set of calibrated parameter sets can be found in the supplementary material.

| Parameter | Initial Range | Units | Refs |
|---|---|---|---|
| **Bacteria** | | | |
| *mtbInternalDoublingTime* | 23,69 | Hours | (50) |
| *mtbExternalDoublingTime* | 23,69 | Hours | (50) |
| **Macrophages** | | | |
| *activatedMacrophageProportion* | 0,0.1 | Per tick | e |
| *baseKillingProbability* | 0.0001,0.02 | Per tick | e |
| *activeKillingProbability* | 0.002,0.3 | Per tick | e |
| *basePhagocytosisProbability* | 0,1 | Per tick | f |
| *activePhagocytosisProbability* | 0,1 | Per tick | f |
| *phagocytosisThreshold* | 8,12 | Internal bacteria | (51) |
| *cellularDysfunctionThreshold* | 8,12 | Internal bacteria | (51) |
| *nfkbSpan* | 0.16,166 | Hours | (31) |
| *TNFthresholdForNFkBActivation* | 40,500 | Molecules | e |
| *bacThresholdForNFkBActivation* | 20,150 | External bacteria | (51) |
| *stat1Span* | 0.16,166 | Hours | (31) |
| *IFNthresholdForStat1Activation* | 40,500 | Molecules | e |
| *ActivatedMacrophageTNFSecretion* | 0,40 | Molecules/second | (52) |
| *InfectedMacrophageTNFSecretion* | 0,40 | Molecules/second | (52) |
| *macrophagePopulation_MaxLifespan* | 20,100 | Days | (51) |
| *macrophagePopulation_MaxActivatedLifespan* | 7,13 | Days | (51) |
| *baseMovementProbabilityMacro* | 0.5,1 | Per tick | (53–56) |
| *activatedMovementProbabilityMacro* | 0,0.5 | Per tick | e |
| **CD4+ T cells** | | | |
| *fractionCD4* | 0.5,0.65 | CD4+ T cells/ CD3+ T cells | (57,58) |
| *fractionTBSpecific* | 0.0001,0.06 | TB specific CD4+ T cells/ Total CD4+ T cells | (59,60) |

| | | | |
|---|---|---|---|
| *activatedTBSpecificCD4Fraction* | 0,0.1 | Initial activated TB specific CD4 T cells/ Total TB specific CD4 T cells | e |
| *CD4ActivationProbability* | 0,1 | Per tick | f |
| *CD4DeactivationProbability* | 0,1 | Per tick | f |
| *ActivatedCD4TNFSecretion* | 0,40 | Molecules/second | (52) |
| *ActivatedCD4IFNSecretion* | 0,40 | Molecules/second | (52) |
| *cd4PopulationDoublingTime* | 6,16 | Hours | (61,62) |
| *maximumCD4Generations* | 3,10 | Generations | (61,63,64) |
| *cd4Population_MaxLifespan* <br> *cd8Population_MaxLifespan* | 34,340 | Days | (65–67) |
| *cd4Population_ActivatedLifespan* <br> *cd8Population_MaxActivatedLifespan* | 2.5,4 | Days | (51,61) |
| *baseMovementProbabilityCD4* <br> *baseMovementProbabilityCD8* | 0,1 | Per tick | f |
| *activatedMovementProbabilityCD4* <br> *activatedMovementProbabilityCD8* | 0,1 | Per tick | f |
| **CD8+ T cells** | | | |
| *CD8Fraction* | 0.3,0.35 | CD8+ T cells/ CD3+ T cells | (58) |
| *tbSpecificCD8Fraction* | 0.0001,0.06 | TB specific CD8+ T cells/ Total CD8+ T cells | (59,60) |
| *activatedTBSpecificCD8Fraction* | 0,0.1 | Initial activated TB specific CD8 T cells/ Total TB specific CD8 T cells | e |
| *CD8ActivationProbability* | 0,1 | Per tick | f |
| *CD8DeactivationProbability* | 0,1 | Per tick | f |
| *ActivatedCD8TNFSecretion* | 0,40 | Molecules/second | (52) |
| *ActivatedCD8IFNSecretion* | 0,40 | Molecules/second | (52) |
| *cd8PopulationDoublingTime* | 3,13 | Hours | (62) |
| *maximumCD8Generations* | 7,20 | Generations | (63,64,68) |
| *CD8KillProbability* | 0.012,0.12 | Per tick | (51) |

| Diffusion | | | |
| --- | --- | --- | --- |
| *TNFthresholdForImmuneCellMovement* | 1,500 | Molecules | e |
| *TNFDiffusionCoefficient* | 0.1,1 | 10^-7 cm^2/s | (31) |
| *TNFDegradationRatePerSecond* | 0.96,10 | 1/s | e |
| *IFNDiffusionCoefficient* | 0.1,1 | 10^-7 cm^2/s | (31) |
| *IFNDegradationRatePerSecond* | 0.96,10 | 1/s | e |
| *granulomaFractionOfDiffusion* | 0,1 | - | f |
| *sphereEfficiency* | 0.65,0.9 | - | e |

**Table 2**: Parameters that were held constant during sampling, their values, and units.

| Parameter | Constant value | Units |
|---|---|---|
| **Traditional v Spheroid** | | |
| CaseNumber | Spheroid: 9<br>Traditional: 15 | - |
| isSpheroid | Spheroid: 1<br>Traditional: 0 | - |
| gridDim_X_Y | Spheroid: 80<br>Traditional: 216 | Grid squares |
| gridDim_Z | Spheroid: 80<br>Traditional: 11 | Grid squares |
| **Simulation defined** | | |
| isBatchRun | 1 | - |
| cellsNeededToBeAddedToGran | 8 | Neighboring immune Cells |
| isPlainColors | 6 | - |
| granQualificationImmuneCellCount | 27 | Immune cells |
| randomSeed | Random | - |
| DiffusionTimeStepMultiplier | 4 | - |
| timestep | 6 | Minutes/step |
| agentLimit | 60000000 | Agents |
| cellsNeededToRemainInGran | 8 | Neighboring immune cells |
| divisionBiomassThreshold | 2 | - |
| **Experimentally defined** | | |
| daysToRun | 6 | Days |
| fractionCD3 | 1 | CD3+ cells/lymphocytes |
| InitialPBMCs | 100000 | Cells |
| fractionMonocyte | 0.4 | Monocyte/PBMC |
| fractionLymphocyte | 0.6 | Lympocyte/PBMC |
| timeToAddTcells | 48 | Hours |
| **Variance** | | |
| cd4DoublingTimeVariance<br>cd8DoublingTimeVariance | 0.25 | |
| nfkbVariance | 0.1 | |
| mtbGrowthRateVariance | 0.1 | |
| stat1Variance | 0.1 | |
| macrophageLifeSpanVariance | 0.1 | |
| cd4LifeSpanVariance<br>cd8LifeSpanVariance | 0.1 | |
| divisionBiomassVariance | 0.2 | |
| newMtbPlacementRange | 0.2 | |
| MinBurstLimit | 20 | Internal bacteria (69) |
| MaxBurstLimit | 40 | Internal bacteria (69) |

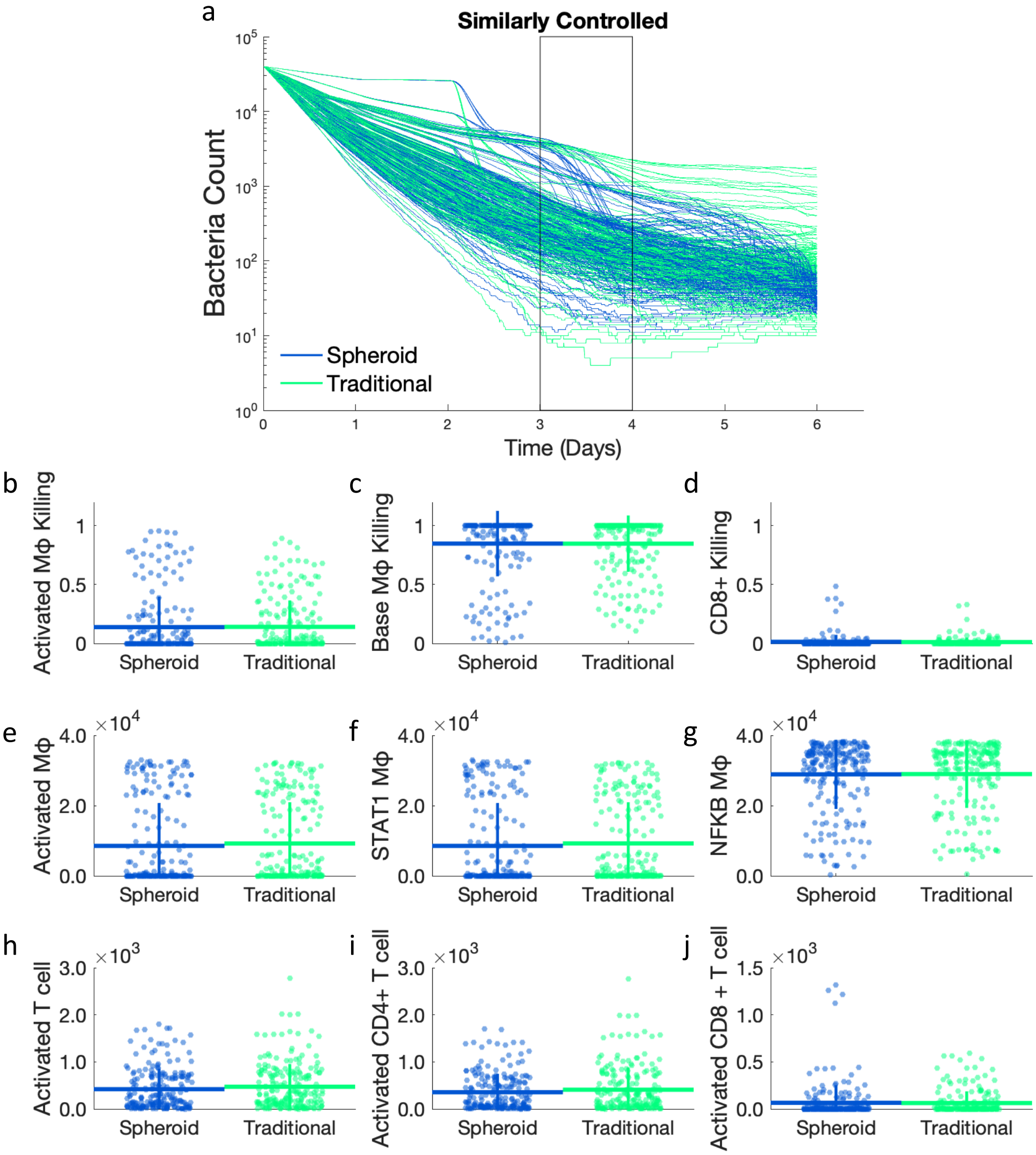


**Figure 2:** a) Bacterial dynamics for similarly controlled simulations. The time range of interest (Day 3-4) is outlined. These are the same time courses as shown in Figure 3b. Proportions of bacterial killing due to b) activated macrophages, c) base macrophages, and d) CD8+ T cell killing from day 3 to 4. Comparing total counts of e) activated macrophages, f) STAT1 activated macrophages, g) NF-κB activated macrophages, h) activated T cells, i) activated CD4+ T cells, and j) activated CD8+ T cells between spheroid and traditional simulations at day 4.

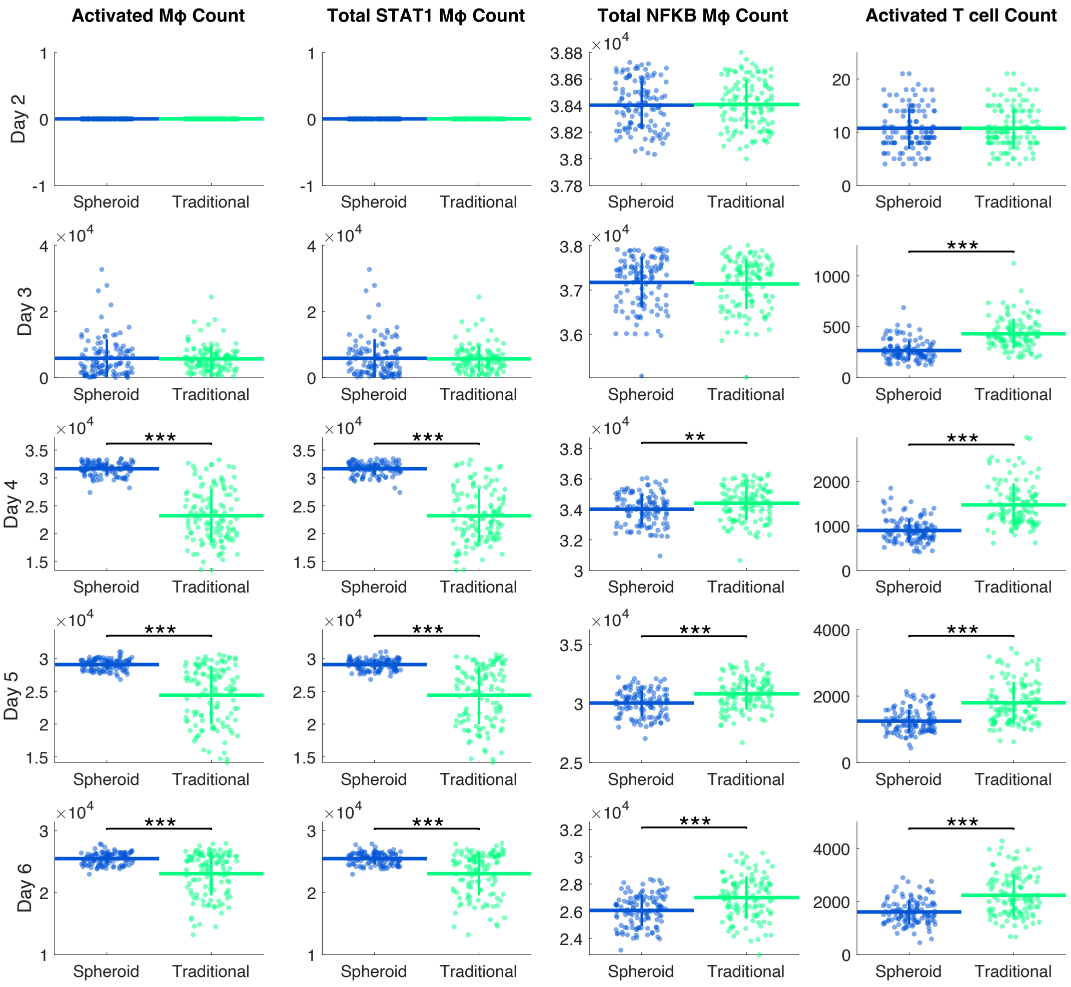


**Figure 3:** Number of activated macrophages, STAT1 macrophages, NFκB macrophages, and T cells from day 2 to day 6 in DC-1 simulations. ** p<=1e-2, *** p<=1e-3

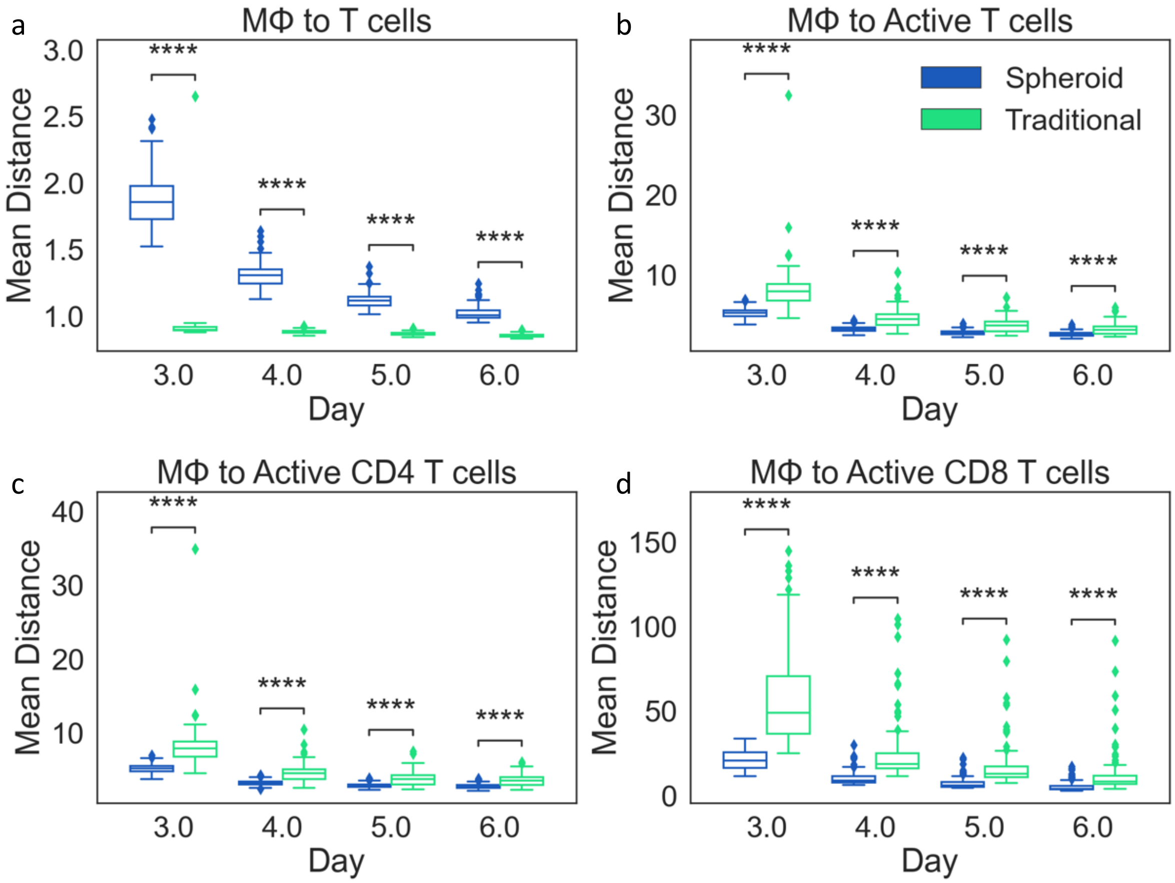


**Figure 4:** The mean distance from macrophages to nearest a) T cells, b) activated T cells, c) activated CD4+ T cells, and activated CD8+ T cells. Each data point represents an average of all cells in a single simulation. The distribution of the mean distances across the set of simulations is compared between DC-1 spheroid and traditional simulations from day 3 to day 6.

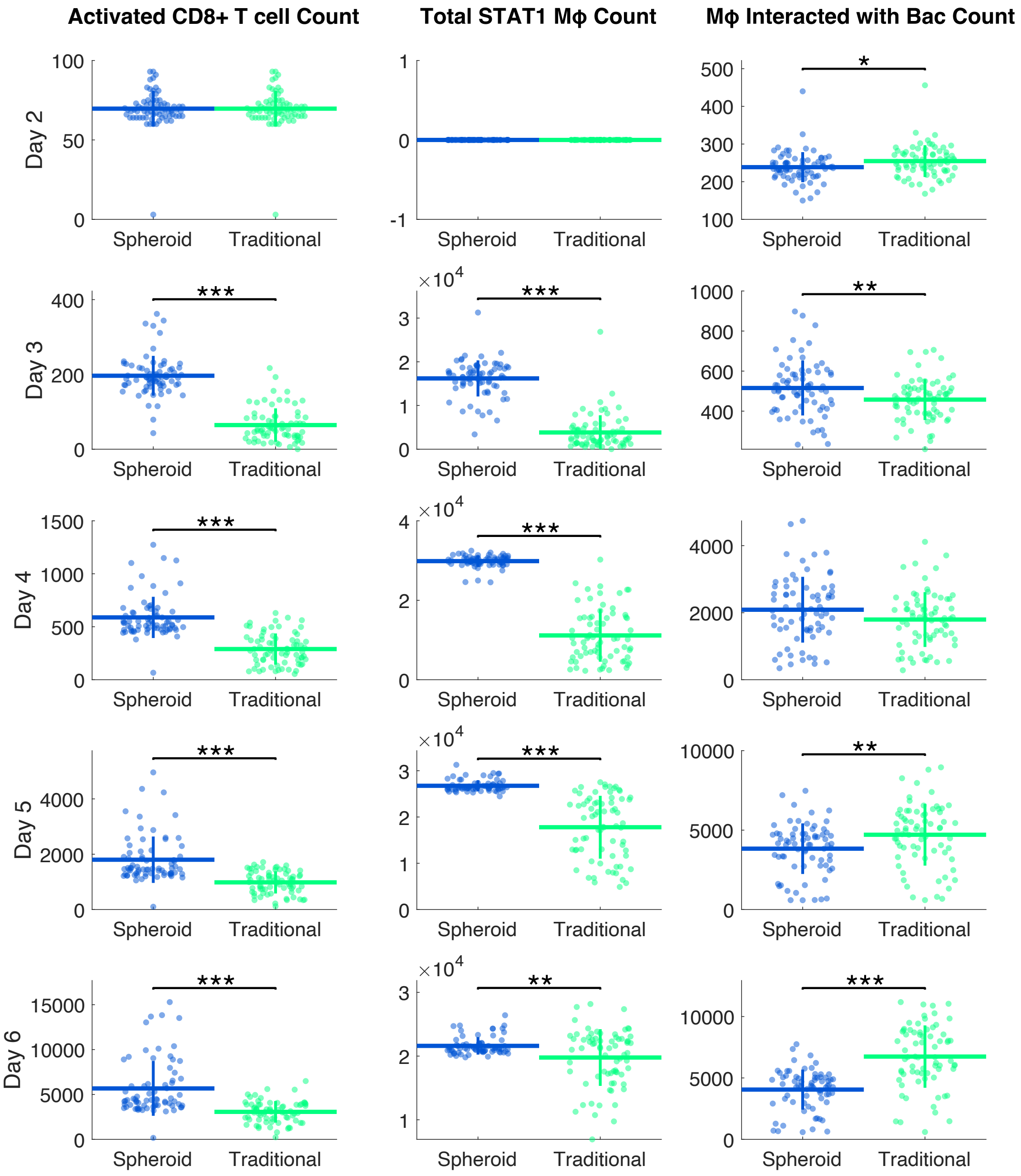


**Figure 5:** Number of activated CD8+ T cells, STAT1 macrophages, and macrophages that have interacted with bacteria from day 2 to day 6 in DC-2 simulations. * p<=1e-1, ** p<=1e-2, *** p<=1e-3

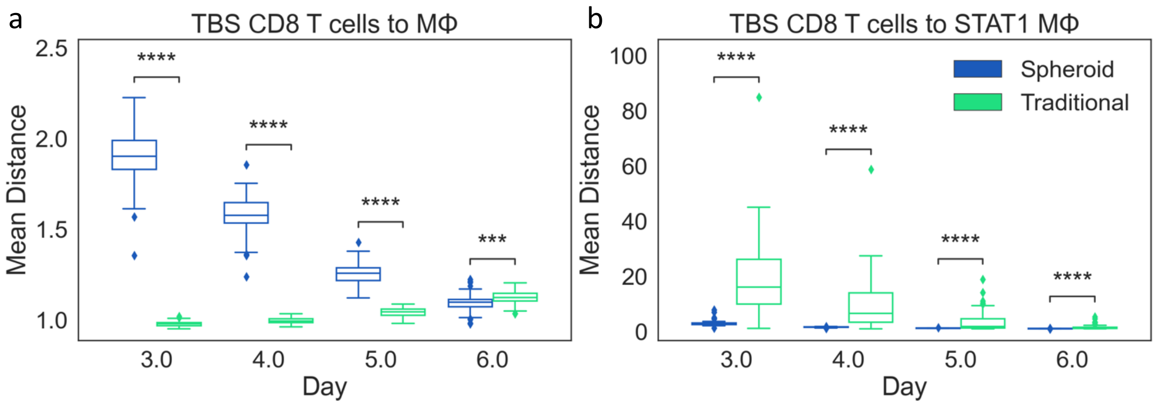


**Figure 6:** The mean distance from a) TB-specific CD8+ T cells to nearest macrophage and b) TB-specific CD8+ T cells to nearest STAT1 activated macrophage. Each data point represents an average of all cells in a single simulation. The distribution of the mean distances across the set of simulations is compared between DC-2 spheroid and traditional simulations from day 3 to day 6. * $p \le 0.05$, *** $p \le 1e-3$, **** $p \le 1e-4$

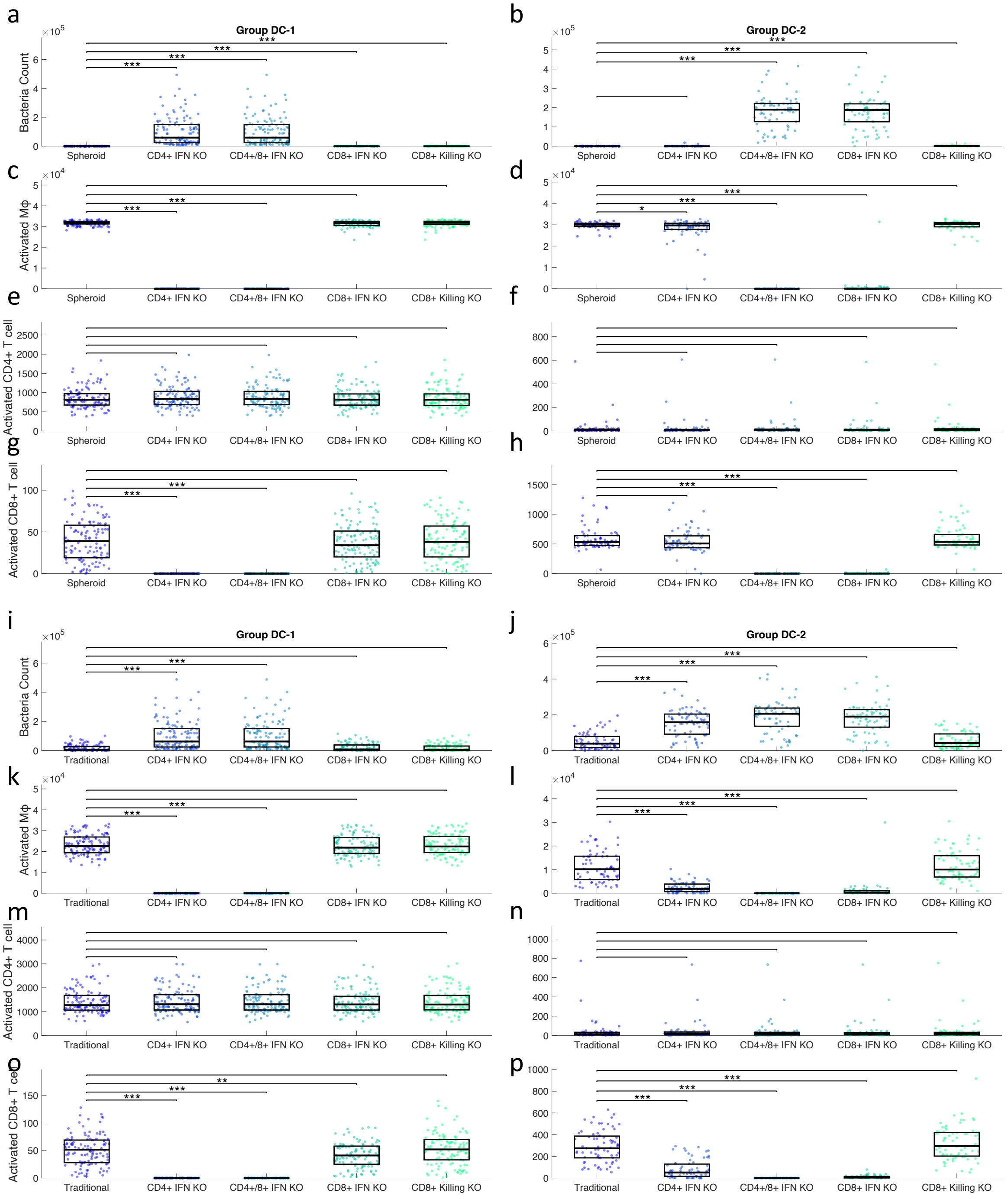


**Figure 7:** Simulations were run with IFN-γ secretion knockouts from CD4+ T cells, CD8+ T cells, or both. A simulation knocking out CD8+ T cell killing of infected macrophages was also performed. The original simulation and the 4 KO scenarios were compared by looking at ab,I,j) bacterial counts at day 6, c,d,k,l) activated macrophage counts at day 4, e,f,m,n) activated CD4+ T cells counts at day 4, and g,h,o,p) activated CD8+ T cell counts at day 4. These outputs are shown for the DC-1 spheroid simulations (a,c,e,g), DC-2 spheroid simulations (b,d,f,h), DC-1 traditional simulations (I,k,m,o), and DC-2 traditional simulations (j,l,n,p).

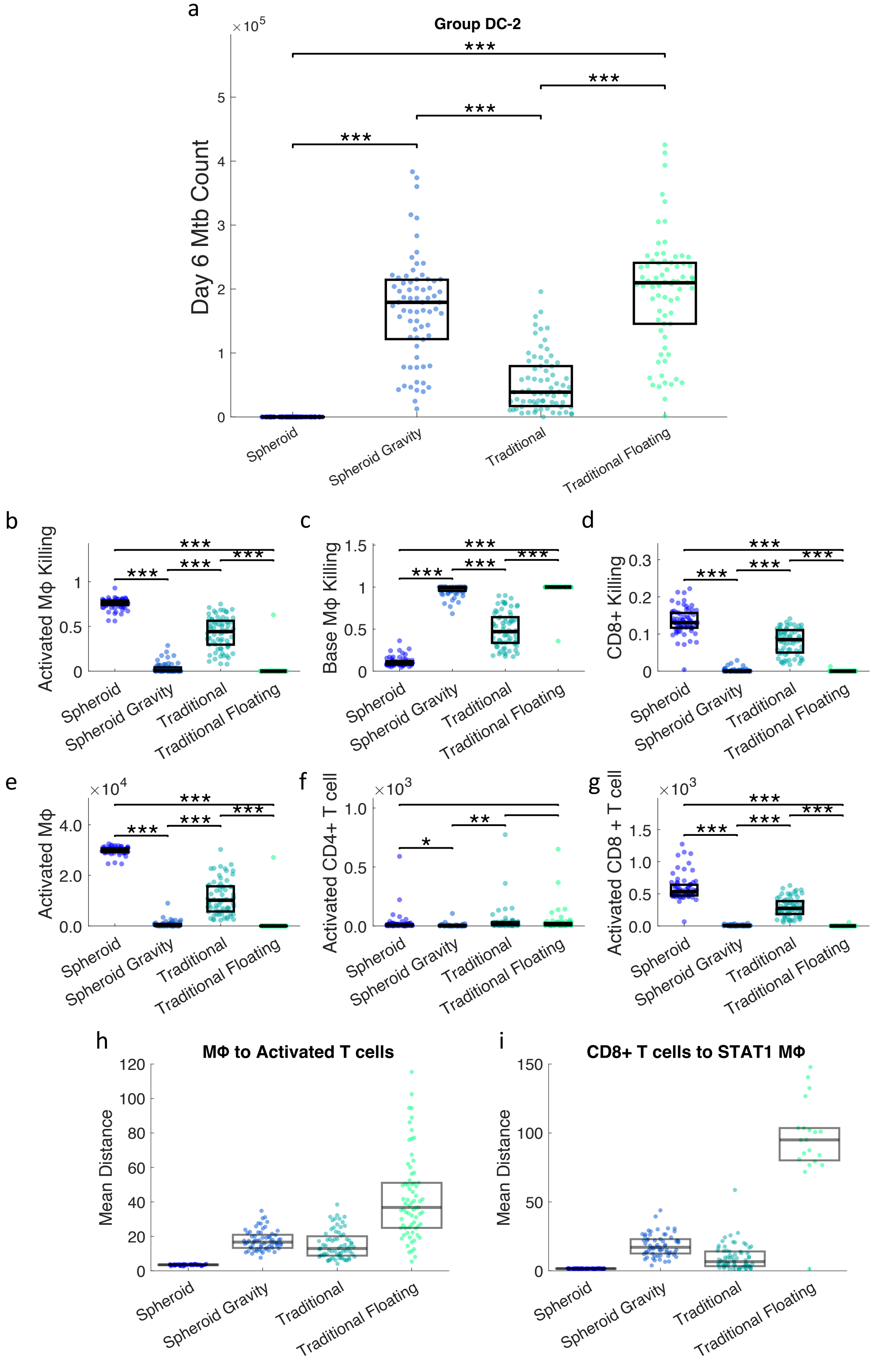
a
Group DC-2
Day 6 Mtb Count
×10^5
***
Spheroid
Spheroid Gravity
Traditional
Traditional Floating
b
Activated MΦ Killing
c
Base MΦ Killing
d
CD8+ Killing
e
Activated MΦ
×10^4
f
Activated CD4+ T cell
×10^3
g
Activated CD8 + T cell
×10^3
h
MΦ to Activated T cells
Mean Distance
i
CD8+ T cells to STAT1 MΦ
Mean Distance

**Figure 8:** DC-2 simulations movement rules swap experiment results. Comparisons between the original setups (spheroid and traditional) and those with swapped movement rules (spheroid gravity and traditional floating) for *Mtb* count at day 6(a); proportion of bacterial killing due to activated macrophages (b), base macrophages(c), and CD8+ T cell cytotoxic killing of macrophages and internalized bacteria(d) from day 3-4; and levels of activation in macrophages(e), CD4+ T cells(f), and CD8+ T cells(g). * p<= 0.05, ** p<=1e-2, *** p<=1e-3

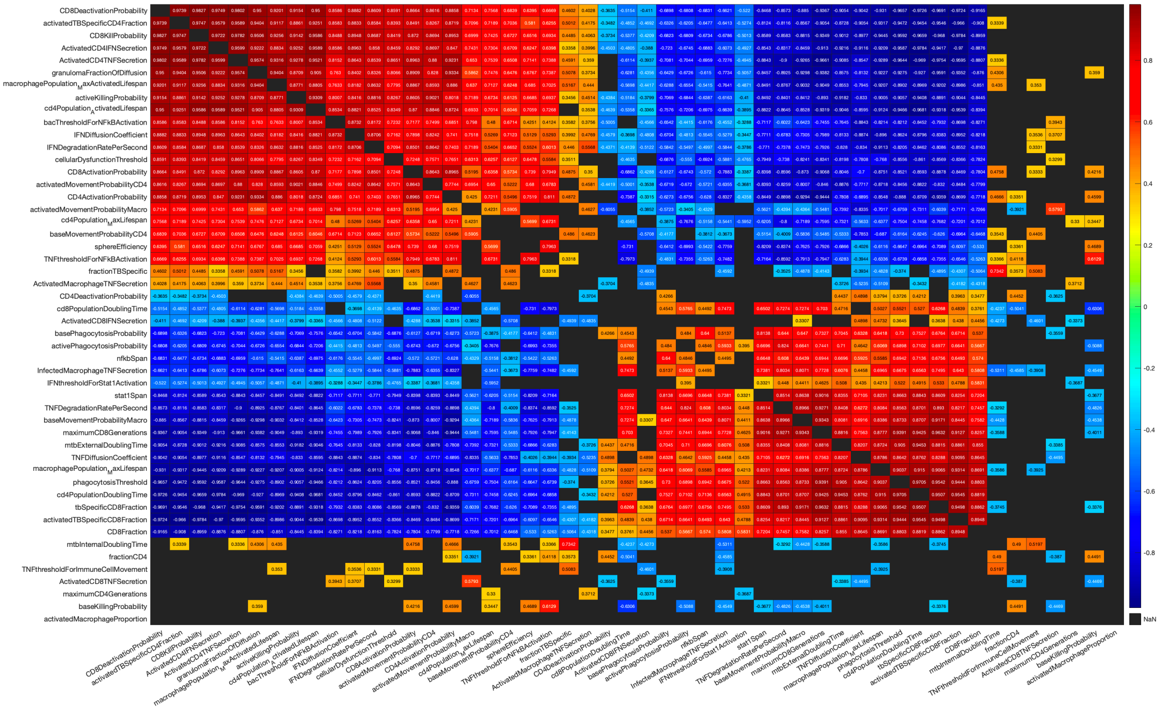

**Figure 9:** Covariance of varied parameters for DC-1 and DC-2 simulations. (p= 0.01 with Bonferroni correction)
